# The Anti-Cancer Effects of Selected Indigenous Medicinal Plants of the Arid Bioregion

**DOI:** 10.64898/2026.08.27.746100

**Authors:** Felix Wambua Muema, Suzanne Thompson, Gerry Turpin, Graham Ambridge, Joanne Jamie, Darren Crayn, Catherine M. Miller, Lionel Hebbard, Phurpa Wangchuk

## Abstract

**Ethnopharmacological relevance:** Australian Indigenous medicinal plants represent a valuable yet underexplored source of bioactive compounds with potential therapeutic relevance. The Iningai community of Central Queensland has traditionally used native plants to manage conditions associated with inflammation, pain, infection, and general illness. Scientific evaluation of these plants may provide evidence for their customary applications and identify bioactivities relevant to anticancer biodiscovery.

**Aim of the study:** This study evaluated leaf and stem extracts of seven medicinal plants— *Pittosporum angustifolium, Alphitonia excelsa, Calytrix microcoma, Geijera parviflora, Melaleuca uncinata, Gossypium australe*, and *Eucalyptus similis*—traditionally used by the Iningai community, focusing on three biological processes relevant to cancer: oxidative stress, inflammation, and cellular proliferation.

**Materials and methods:** Antioxidant activity was assessed using DPPH radical-scavenging and ferric reducing antioxidant power (FRAP) assays. Anti-inflammatory activity was evaluated in lipopolysaccharide (LPS)-stimulated THP-1 macrophage-like cells by profiling IFN-α, TNF-α, IL-6, IL-12, IL-18, and IL-23. Antiproliferative activity was assessed using MTT-based viability assays in human and murine liver cancer cell lines (Huh7, Hep3B, Hep-55.1c, and A52).

**Results:** The extracts exhibited distinct biological activity profiles. *G. parviflora* stem and *C. microcoma* leaf extracts showed the strongest antioxidant activities, whereas *P. angustifolium* stem exhibited the weakest radical-scavenging capacity. Cytokine responses were extract-specific, with *E. similis* leaf extract demonstrating broad and pronounced suppression of multiple LPS-induced pro-inflammatory cytokines. Several extracts produced concentrationdependent reductions in liver cancer cell viability, with *P. angustifolium* stem exhibiting the most consistent and potent antiproliferative activity across the cell lines tested. Notably, strong antioxidant or anti-inflammatory activity did not necessarily correspond with antiproliferative activity.

**Conclusion:** Australian Indigenous medicinal plant extracts demonstrated distinct antioxidant, immunomodulatory, and antiproliferative activities rather than uniform bioactivity across experimental systems. The divergent activities of *G. parviflora, C. microcoma, E. similis*, and *P. angustifolium* highlight the importance of integrated biological screening and support the value of Indigenous knowledge-guided biodiscovery. These plants represent promising sources for further investigation of selective bioactive compounds with potential relevance to anticancer drug discovery.

## 1.0 Introduction

Chronic inflammation and oxidative stress are closely interconnected biological processes that play central roles in the pathogenesis of numerous diseases, including cancer, metabolic disorders, and cardiovascular conditions (Liu et al., 2025; Olivares-Vicente and Herranz-López, 2025). Excessive production of reactive oxygen species (ROS) can disrupt cellular redox homeostasis, leading to oxidative damage to lipids, proteins, and DNA (Jomova et al., 2023). In parallel, sustained inflammatory signalling, often mediated by pro-inflammatory cytokines and activated immune cells, contributes to tissue damage and disease progression (Zhao et al., 2021). Consequently, agents capable of simultaneously modulating oxidative stress and inflammatory responses are of significant therapeutic interest.

Macrophages are key regulators of innate immunity and inflammation (Mussbacher et al., 2023). Upon exposure to bacterial components such as lipopolysaccharide (LPS), macrophages adopt a pro-inflammatory phenotype characterized by elevated production of cytokines and inflammatory mediators. While this response is essential for host defence, its dysregulation is associated with chronic inflammatory diseases and cancer development (Mussbacher et al., 2023; Ni et al., 2023). In the tumour microenvironment, inflammatory mediators can promote cancer cell survival, proliferation, angiogenesis, and immune evasion (Jiang et al., 2020). Therefore, the identification of natural compounds that can attenuate LPS-induced inflammatory responses, while exerting selective cytotoxicity toward cancer cells, represent a promising strategy for disease prevention and therapy.

In addition to their roles in oxidative stress and inflammation, dysregulated cellular proliferation is a defining hallmark of cancer progression (Wang et al., 2025). Of relevance to our work, numerous studies have shown that specific inflammatory factors, for example IFN-α, IL-12 and IL-18 can restrict, while IL-6, TNFα and IL-23 can promote human hepatocellular carcinoma (HCC) growth and progression (Grivennikov et al., 2010; Naugler et al., 2007). In this light, natural products have emerged as important sources of antiproliferative agents, with numerous plant-derived compounds demonstrating the ability to inhibit cancer cell growth through modulation of inflammatory driven pathways. For example, flavonoids such as quercetin and kaempferol can suppress proliferation and induce apoptosis in liver cancer cells (Li et al., 2025; Singh et al., 2025), while alkaloids like berberine exhibit growth-inhibitory effects through disruption of mitochondrial function and cell cycle arrest (Almatroodi et al., 2022). These highlight the therapeutic potential of plant-derived metabolites not only as antioxidants or anti-inflammatory agents, but also as direct modulators of cancer cell proliferation.

Australia possesses uniquely diverse flora (Andrew et al., 2021), with many native plants used for millennia in Indigenous medicinal practices (Turpin et al., 2022). Australian Indigenous medicinal plants, particularly arid plants, represent an underexplored reservoir of bioactive compounds with potential therapeutic applications (Turpin et al., 2022). Traditional knowledge has long recognised the use of these plants for treating inflammation, infections, wounds, and other ailments, yet scientific validation of their biological activities remains limited (Turpin et al., 2025; Yeshi et al., 2022). Systematic investigation of these plants, through partnerships between Traditional Owners and Western scientists where access and benefit sharing agreements are negotiated with the full prior informed consent of knowledge holders, may not only support the preservation and recognition of Indigenous knowledges but also contribute to the discovery of novel bioactive agents.

In recent years, growing interest has emerged in evaluating Australian native plant extracts for their antioxidant, anti-inflammatory, and anticancer properties (Turpin et al., 2022). Nevertheless, comprehensive studies that integrate antioxidant profiling with immunomodulatory effects in macrophage models and cytotoxicity assessment in cancer cell lines are scarce. In particular, understanding whether plant extracts suppress inflammatory responses or, conversely, exacerbate cytokine production is critical for accurately interpreting their therapeutic relevance.

This study investigated the biological activities of selected Australian Indigenous medicinal plant extracts of Iningai Country located in Central Western Queensland, Australia, extending across the region surrounding Barcaldine and towards Longreach, Aramac and Muttaburra. The region forms part of Australia’s semi-arid region and encompasses landscapes characterised by open grasslands and woodland communities adapted to highly variable rainfall and periodic drought. These environmental conditions contribute to a distinctive native flora adapted to climatic and ecological stress. The Iningai people have maintained longstanding cultural relationships with this landscape and its plant resources, including the use of native species for medicinal purposes. This traditional knowledge provides an important basis for investigating the pharmacological properties of plants occurring on Iningai Country. The anti-cancer activities were evaluated on human and murine liver cancer cell lines, by evaluating their antioxidant, immunomodulatory and antiproliferative effects. This study is part of the pathway of their scientific examination and potential therapeutic development.

## 2.0 Materials and Methods

### 2.1 Plant materials collection

The plant materials (*Pittosporum angustifolium, Alphitonia excelsa, Calytrix microcoma, Geijera parviflora, Melaleuca uncinata, Gossypium australe*, and *Eucalyptus similis*) were obtained from Turraburra Station on Iningai Country in Central Western Queensland, Australia, through the Yumbangku Aboriginal Cultural Heritage and Tourism Development Aboriginal Corporation (YACHATDAC), Barcaldine, Queensland, Australia. Plant selection was informed by traditional medicinal knowledge shared by the Iningai community. Plant identification was verified by a senior ethnobotanist, Gerry Turpin and staff of the Queensland Herbarium (Index Herbariorum code BRI). Voucher specimens were deposited at the Australian Tropical Herbarium (Index Herbariorum code CNS).

### 2.2 Sample preparation and crude extraction

The leaves were separated from the stems, thoroughly washed with water, and oven-dried at 40 °C. The leaves of *A. excelsa, C. microcoma*, and *E. similis*, stems of *P. angustifolium, G. parviflora*, and *G. australe*, and twigs of *M. uncinata* were then ground into a coarse powder using a NutriBullet® blender followed by ultra-sonic assisted crude extraction (Niu et al., 2025), using 20 g of sample resuspended in ethanol (80%). The ethanol crude extraction process was repeated three times with fresh solvent each time. The extracts were combined and filtered with a Stericup system (0.22 μm PES, Merck, USA), the solvent eliminated under reduced pressure, at 40 °C using a rotary evaporator (Heidolph, Germany) and the extract stored at −30 °C.

### 2.3 DPPH Radical Scavenging Assay

The free radical scavenging activity of the plant extracts was evaluated using the 2,2-diphenyl-1-picrylhydrazyl (DPPH) assay (Muema et al., 2022). Briefly, freshly prepared DPPH solution (0.1 mM) in methanol was mixed with concentrations (7.8125 - 500 µg/mL) of the crude plant extracts. 100 µL of test sample (prepared crude extract samples and standard) was combined with 100 µL of DPPH solution in a 96-well flat-bottom microplate. The reaction mixtures were shaken and incubated in the dark at room temperature for 30 min, after which the absorbance was measured at 517 nm using a microplate reader (SPECTROstar® Omega, BMG Labtech, Mornington, Australia). Methanol was the blank, DPPH solution without extract was the negative control, and vitamin C (ascorbic acid) was the reference antioxidant standard. The percentage of DPPH radical scavenging activity was calculated: Scavenging activity (%) = (A_c_ - A_s_/ A_c_) × 100; where, A_c_ and A_s_ are the absorbance of the control and the reaction sample, respectively. IC□ □was calculated using GraphPad Prism software (version 10.0.2) and expressed in µg/mL.

### 2.4 Ferric Reducing Antioxidant Power (FRAP) Assay

The ferric reducing antioxidant power (FRAP) assay was performed (Mutungi et al., 2021) to determine the reducing capacity of the extracts. The FRAP reagent was freshly prepared by mixing acetate buffer (300 mM, pH 3.6), 2,4,6-Tri(2-pyridyl)-*s*-triazine (TPTZ) solution (10 mM in 40 mM HCl), and FeCl□·6H□O (20 mM) in a 10:1:1 (v/v/v) ratio. 10 μL of each extract/standard was added to 190 μL of FRAP working solution in a 96-well microplate and incubated at 37 °C for 20 min. Absorbance was measured at 593 nm and a standard calibration curve was generated using ferrous sulphate (FeSO_4_·7H_2_O, 6.25–2000 mmol). Antioxidant capacity was expressed as mmol Fe^2+^/g of the sample.

### 2.5 Cell Culture

Ethical approval for this assay was obtained from the James Cook University Human Research Ethics Committee under approval number 22H-8072. Human monocytic leukemia cells THP-1 (ATCC TIB-202) were cultured in RPMI-1640 medium, 10% foetal bovine serum (FBS) and 1% penicillin-streptomycin (P/S). Human hepatocellular carcinoma (HCC) cell lines Huh7 and Hep3B, and mouse HCC Hep-55.1c, and A52 cells were cultured in DMEM 10% FBS and 1% P/S and incubated at 37 °C in a humidified atmosphere containing 5% CO□.

### 2.6 Cell Viability Assay (MTT)

Viability of the THP-1 cell was assessed with the MTT assay. Cells were differentiated into macrophage-like cells using phorbol 12-myristate 13-acetate (PMA, 100 ng/mL), seeded into 96-well U-shaped culture plates (Falcon®, Corning, NY, USA) at a density of 1 × 10^4^ cells per well (100 μL) and cultured overnight at 37 °C in a humidified atmosphere containing 5% CO□. The cells were then treated with the plant extracts at concentrations of 100, 75, 50, 25, and 10 μg/mL (dissolved in RPMI-1640 media). Every 24 hrs, the medium with extract was aspirated and replaced with fresh medium containing extract for 3 days. Following treatment, 15 μL of MTT solution (0.5 mg/mL) was added to each well and the plates were incubated for 4 h at 37 °C. The formazan crystals were dissolved in 50μL dimethyl sulfoxide (DMSO), and absorbance measured at 570 nm. Cell viability was expressed as a percentage relative to untreated control cells, and DMSO was used as a negative control.

### 2.7 Cytotoxicity Evaluation in Liver Cancer Cell Lines

The antiproliferative effects of the extracts on Huh7, Hep3B, Hep-55.1c, and A52 cells were evaluated by the MTT assay (Muema et al., 2022). Cells were seeded in a 96-well culture plates (6 × 10^3^ cells per well) and left overnight, treated with extracts (12.5 - 400 μg/mL) for 48 h. Following treatment, MTT reagent was added and incubated to allow viable, metabolically active cells to reduce the tetrazolium salt to formazan crystals. The crystals were solubilised in DMSO and absorbance measured at 570 nm. Cell viability was calculated as the percentage of absorbance relative to untreated control cells. Reductions in cell viability were interpreted as indicative of antiproliferative and/or cytotoxic effects of the extracts. Dose-response graphs were generated, and half-maximal inhibitory concentration (IC□ □) values calculated using GraphPad Prism software (version 10.0.2).

### 2.8 LPS-Stimulated Inflammatory Response in THP-1 Cells

THP-1 cells were seeded in 48-well culture plates (1×10^6^ cells/well) and differentiated into macrophages cells using PMA. Groups consisted of controls (cells in media), and LPS stimulated cells, with or without extracts, treated for 72 hours, and the supernatants collected for cytokine analysis.

### 2.9 Quantification of inflammatory cytokines

Cytokine profiling of THP-1 culture supernatants was performed with a customized LegendPlex− multi-analyte flow assay kit (BioLegend®, USA, Cat. No. 740809; Lot No. B471106) targeting 13 human inflammatory cytokines (interleukin-1β (IL-1β), interferon-α (IFN-α), interferon-γ (IFN-γ), tumor necrosis factor (TNF), monocyte chemoattractant protein-1 (MCP-1), IL-6, IL-8, IL-10, IL-12, IL-17A, IL-18, IL-23, and IL-33). Assays were performed as per the manufacturer’s instructions on a BD LSRFortessa X20 flow cytometer (BD Biosciences); data was processed using BioLegend®’s cloud-based analysis software (San Diego, CA, USA), and cytokine concentrations were expressed as pg/mL (mean ± SD).

### 2.10 Statistical Analysis

All experiments were conducted in triplicate and repeated independently at least three times. Data are presented as mean ± SEM. Statistical analyses and graphical representations were performed using GraphPad Prism software. One-way analysis of variance (ANOVA) followed by appropriate post hoc tests was used to assess differences between groups. For antioxidant (DPPH) and anticancer (MTT) assays, IC□ □values were calculated using nonlinear regression analysis (dose-response curve fitting) in GraphPad Prism. Differences were considered statistically significant at *p* < 0.05.

## 3.0 Results

### 3.1 Medicinal plants diversity of Iningai and Ethnobotanical uses of seven selected medicinal plants

The medicinal flora used by the Iningai people reflects a diverse assemblage of native Australian plant species spanning multiple taxonomic families, including Pittosporaceae, Rhamnaceae, Myrtaceae, Rutaceae, and Malvaceae. The selected species, *Pittosporum angustifolium, Alphitonia excelsa, Calytrix microcoma, Geijera parviflora, Melaleuca uncinata, Gossypium australe*, and *Eucalyptus similis* (**Table 1 and Figure 1**) demonstrate both botanical diversity and functional convergence in traditional therapeutic applications (Turpin et al., 2026). Notably, leaves and stems constitute the most frequently utilised plant parts, suggesting ease of accessibility and sustainable harvesting practices within customary knowledge systems.

**Table 1.** Ethnobotanical uses of selected medicinal plants from the Iningai region.

| Botanical Name | Common name | Parts collected for studies | Customary uses |
| --- | --- | --- | --- |
| *Pittosporum angustifolium* Lodd., G.Lodd. & W.Lodd. (Pittosporaceae) | Gumbi gumbi | Stem | Used to treat cough and cold, skin diseases, cramps, eczema, muscle aches, bruises and to induce lactation in mothers of newborns. |
| *Alphitonia excelsa* (A.Cunn ex Fenzl) Benth. (Rhamnaceae) | Soap tree, soap bush, red ash | Leaf | Leaves are used as a soap, as antiseptic for skin infections, wounds and sores. |
| *Calytrix microcoma* Craven (Myrtaceae) | Pink desert heather | Leaf | Leaves and flowers are used as insect repellent and in the treatment of cuts, wounds and muscle pain. |
| *Geijera parviflora* Lindl. (Rutaceae) | Wilga | Stem | Used for aches and pains both internally and externally. |
| *Melaleuca uncinata* R.Br. (Myrtaceae) | Prickly-leaved paperbark | Twigs | Used for respiratory issues, stomach issues and aches and pains. |
| *Gossypium australe* F.Muell. (Malvaceae) | Desert rose | Stem | Treating coughs and colds, cuts and sores. |
| *Eucalyptus similis* Maiden (Myrtaceae) | Queensland yellowjacket | Leaf | Used for respiratory and stomach issues as well as pain relief and fever reduction |

**Figure 1.**
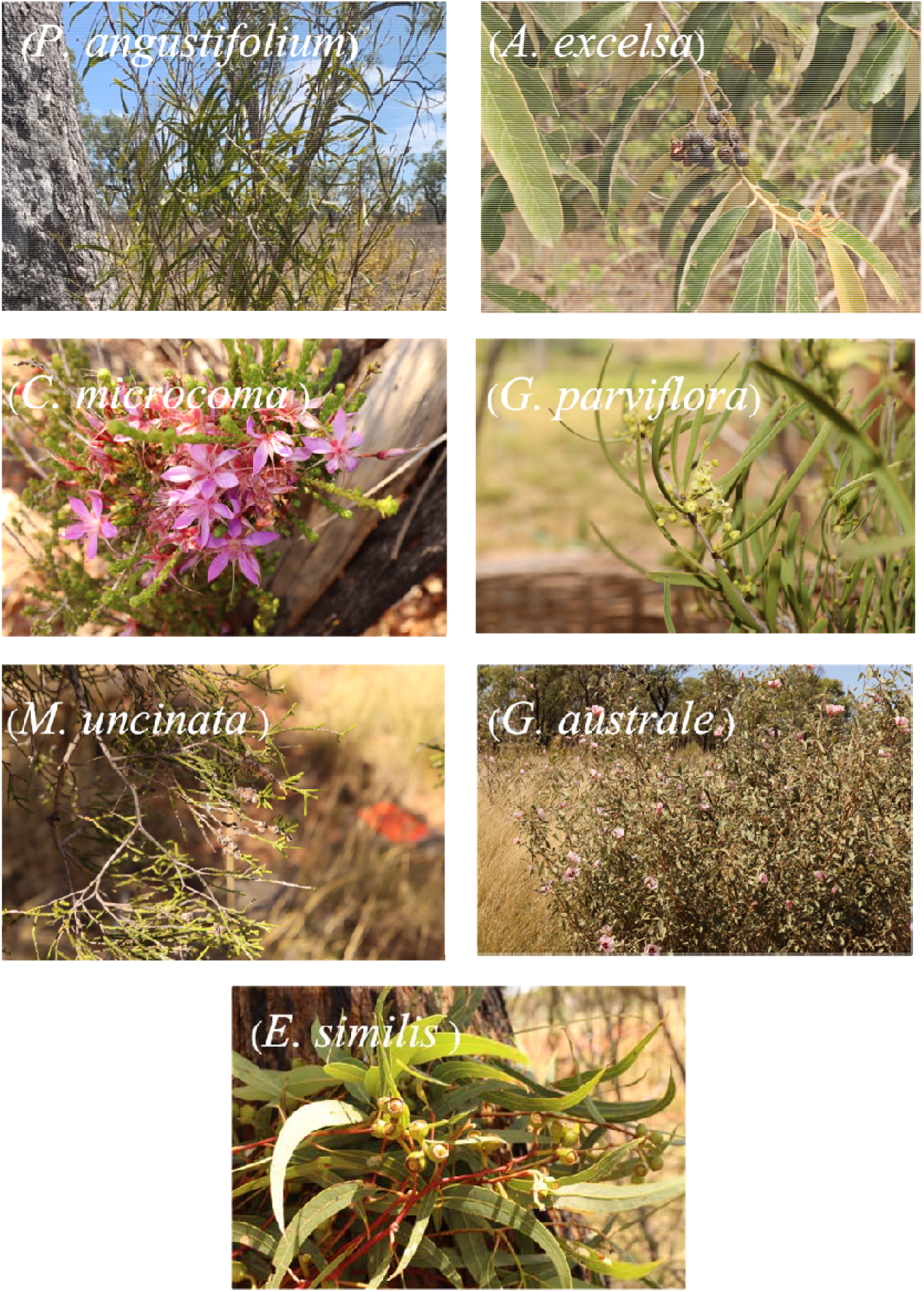
Australian Indigenous medicinal plants included in the study (photos: Gerry Turpin, Lorna Ngugi)

Ethnobotanical records indicate that these plants are predominantly employed in the management of conditions associated with inflammation, infection, and pain. *P. angustifolium* is widely used for respiratory ailments, skin disorders, and musculoskeletal conditions (Blonk and Cock, 2019; Phan et al., 2020), while *A. excelsa* serves as both a cleansing agent and topical antiseptic for wounds and infections (Lavhale and Mishra, 2007). *C. microcoma* and *G. parviflora* are applied in treating cuts, muscle pain, and general aches, highlighting their relevance in anti-inflammatory and analgesic contexts (Lavhale and Mishra, 2007). *M. uncinata* and *E. similis* are traditionally used to address respiratory and gastrointestinal disorders, as well as fever, whereas *G. australe* is employed for treating coughs, colds, and skin injuries.

### 3.2 DPPH radical□scavenging activity and IC□□ values

The free radical scavenging capacity of the seven 80% aqueous extracts was assessed using the DPPH assay and vitamin C (ascorbic acid) as a reference standard. All extracts exhibited DPPH inhibition in a dose-dependent manner (**Figure 2A-B**) and **Table 2** depicts the IC_50_ values of the samples. Among the plant extracts, *G. parviflora* stem exhibited the strongest DPPH radical scavenging activity with an IC□ □value of 8.87 ± 0.3 μg/mL, followed by *C. microcoma* leaf (12.95 ± 1.3 μg/mL), whereas *P. angustifolium* stem displayed the weakest scavenging capacity (53.38 ± 3.7 μg/mL). As expected, the reference antioxidant vitamin C demonstrated greater activity than all extracts, with an IC□ □value of 2.38 ± 0.01 μg/mL.

**Table 2.**
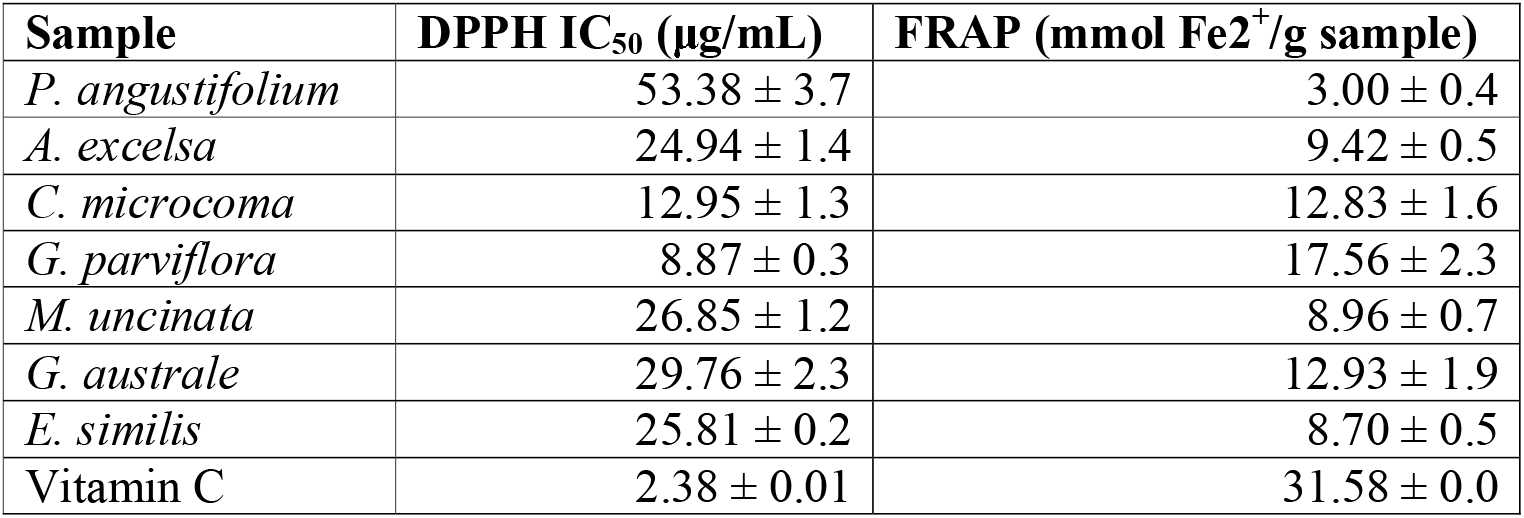
Shows the scavenging capacity and the ferric reducing power of examined plant extracts and the standard.

| Sample | DPPH IC<sub>50</sub> (µg/mL) | FRAP (mmol Fe<sup>2+</sup>/g sample) |
| --- | --- | --- |
| *P. angustifolium* | 53.38 ± 3.7 | 3.00 ± 0.4 |
| *A. excelsa* | 24.94 ± 1.4 | 9.42 ± 0.5 |
| *C. microcoma* | 12.95 ± 1.3 | 12.83 ± 1.6 |
| *G. parviflora* | 8.87 ± 0.3 | 17.56 ± 2.3 |
| *M. uncinata* | 26.85 ± 1.2 | 8.96 ± 0.7 |
| *G. australe* | 29.76 ± 2.3 | 12.93 ± 1.9 |
| *E. similis* | 25.81 ± 0.2 | 8.70 ± 0.5 |
| Vitamin C | 2.38 ± 0.01 | 31.58 ± 0.0 |

**Figure 2.**
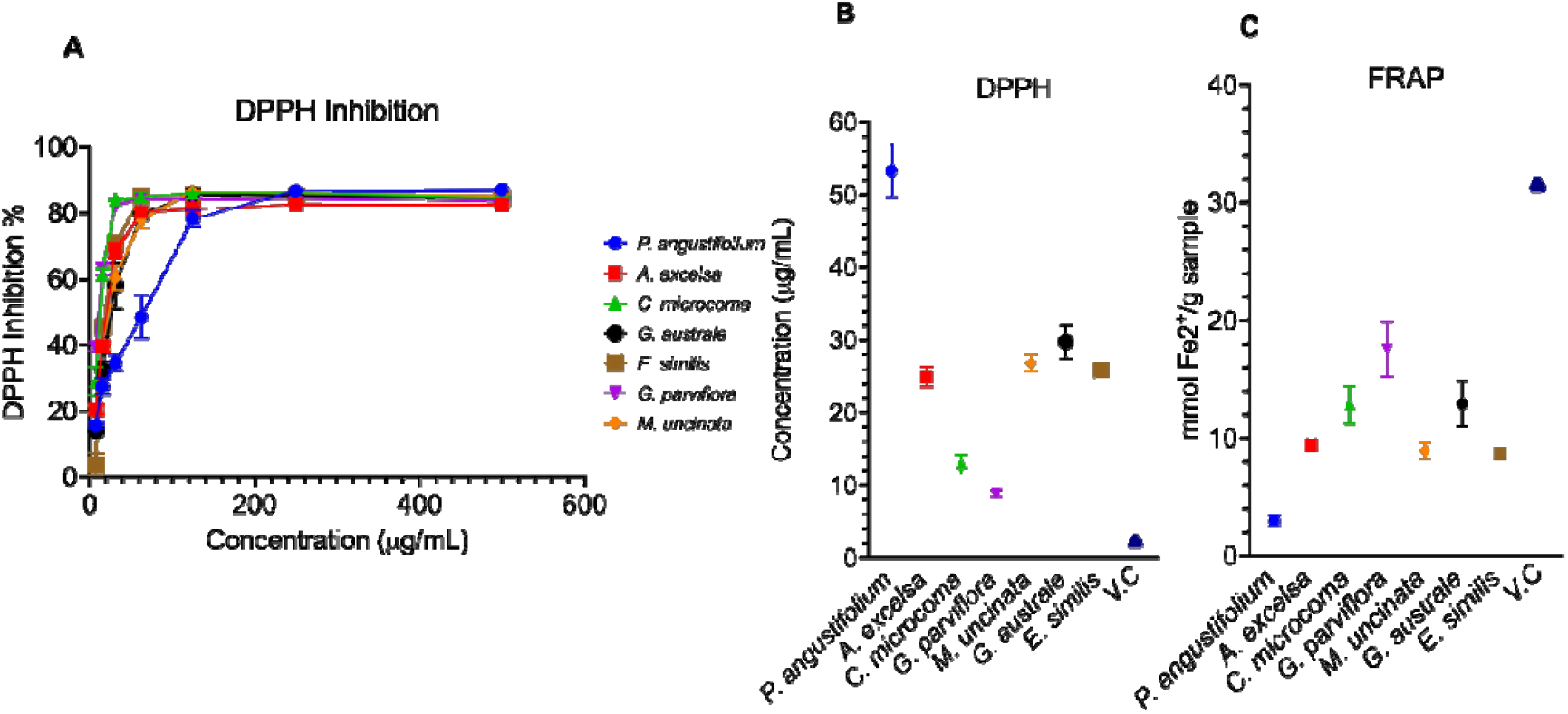
The percentage (%) scavenging rate and antioxidant capacities of the plant extracts. **A**, the dose–response curves for DPPH radical scavenging activity of the extracts over seven concentrations (7.8125–500 µg/mL) while vitamin C (V.C) was examined at concentrations (0.9375-30 µg/mL). **B**, shows the DPPH IC□ □values (µg/mL) for each extract, and vitamin C as a positive control. **C**, depicts the FRAP values for the extracts, expressed as mmol Fe^2^□ /gram of extract. Values represent mean ± SD (n = 3).

### 3.3 Ferric-reducing antioxidant capacity (FRAP)

To confirm antioxidant capacity of the extracts, FRAP studies were performed. Interestingly, there was correlation between the two assays. Among the extracts, *G. parviflora* stem displayed the greatest ferric reducing antioxidant capacity (17.56 ± 2.3 mmol Fe^2^□/g), followed by *G. australe* stem and *C. microcoma* leaf, whereas *P. angustifolium* stem exhibited the lowest activity (3.00 ± 0.4 mmol Fe^2^□/g). The reference standard, vitamin C showed a higher reducing capacity than all extracts (31.58 ± 0.0 mmol Fe^2^□/g). The ferric reducing capacity of all extracts are shown in **Figure 2C** and **Table 2**. Together the results illustrate the specific antioxidant properties of the extracts.

### 3.4 Anti-inflammatory activity

Stimulation with LPS markedly increased cytokine production compared with the untreated controls across all groups (**Figure 3A-F**). The plant extracts were able to modulate cytokine release. *E. similis* significantly reduced IFN-α levels 3.2-fold versus LPS alone (**Figure 3A**). For TNF-α, *P. angustifolium* stem and *M. uncinata* twigsincreased levels 1.7-fold and 1.5-fold, and *A. excelsa, G. australe* and *E. similis*, reduced them 1.9-fold, 3.5-fold and 1.6-fold, respectively, (**Figure 3B**). In evaluating IL-6, *P. angustifolium, M. uncinata* and *G. australe* showed significant increases of 1.5-fold, 2.3-fold and 2.3-fold, respectively, and *E. similis*, significantly suppressed levels by 3.3-fold (**Figure 3C**). For IL-12, a broad inhibitory response was evident in all the extracts, with *C. microcoma* and *E. similis* producing the most substantial decreases of 3.3-fold and 3.2-fold, respectively, compared with LPS alone (**Figure 3D**). In considering IL-18, the extracts *A. excelsa, C. microcoma* and *E. similis* significantly lowered IL-18 levels, by 1.6-fold, 1.7-fold and 2.3-fold, respectively versus control; and *G. australe* significantly increased IL-18 levels by 1.3-fold (**Figure 3E**). For IL-23, *C. microcoma, M. uncinata*, and *E. similis* significantly suppressed IL-23, by 2.0-fold, 3.2-fold and 5.3-fold, respectively, and *A. excelsa* increased IL-23 levels 1.8-fold, relative to the LPS-treated group (**Figure 3F**). These results demonstrate that the plant extracts exert cytokine-specific action on LPS-induced inflammatory responses.

**Figure 3.**
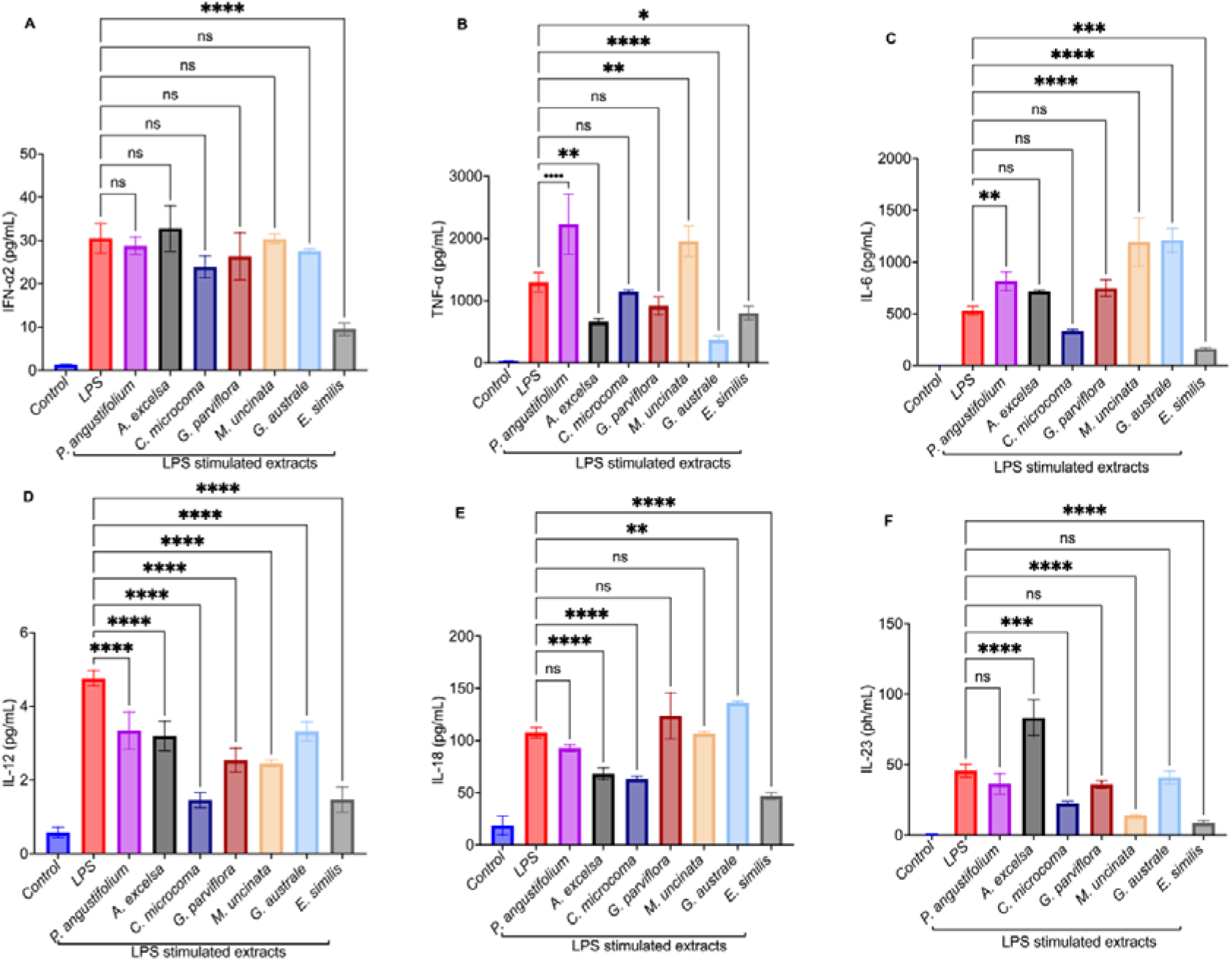
Modulatory effects of the medicinal plant extracts on LPS-induced proinflammatory cytokine production. Graphs show the media concentrations of individual cytokines **(A)** IFN-α, **(B)** TNF-α, **(C)** IL-6, **(D)** IL-12, **(E)** IL-18, and **(F)** IL-23 (pg/ml) of LPS-stimulated cells after treatment with individual plant extracts (*P. angustifolium* – *E. similis*), versus control (media) and LPS-only groups. Bars represent mean ± SD from replicate experiments (n=3), with individual data points shown where applicable. Statistical significance was determined by one-way ANOVA, and significance indicated as, p < 0.05 (*), p < 0.01 (**), p < 0.001 (***), and p < 0.0001 (****).

### 3.5 Anti-cancer activity

The anti-proliferative effects of the plant extracts were evaluated on four cancer cell lines, Huh7, Hep3B, Hep-55.1c, and A52, by the MTT assay, after 48 hrs exposure (12.5–400 µg/mL; **Figure 4**). Only *P. angustifolium, C. microcoma, G. parviflora*, and *M. uncinata* extracts reduced cell viability. For Huh7 cells, these extracts produced a clear concentrationdependent reduction in viability. *P. angustifolium* and *M. uncinata* showed the strongest effects, with marked decreases in viability of 91 and 63%, respectively, at ≥100 µg/mL. *C. microcoma* and *G. parviflora* had moderate inhibitory activity, with 400 µg/mL reducing proliferation by 60 and 39%, respectively. In Hep3B cells, *P. angustifolium* stem demonstrated the most pronounced cytotoxic effect, promoting 80% loss of cell viability at 200 µg/mL. *C. microcoma* reduced viability in a dose-dependent manner, but to a lesser extent than *P. angustifolium*, while at 200 µg/mL, viability was reduced by 60%. In Hep-55.1c cells, the extracts showed concentration-dependent antiproliferative activity. *P. angustifolium* stem produced the strongest reduction in cell viability, followed by *C. microcoma* leaf and *M. uncinata* twigs of 85, 67 and 82%, respectively at 400 µg/mL. In A52 cells, dose-dependent trends were also observed. *P. angustifolium* stem caused the greatest reduction in cell viability across the concentration range, at 100 and 400 µg/mL, proliferation was reduced by 62 and 90%, respectively. Whereas *C. microcoma* and *M. uncinata* showed less inhibitory effects, that only became more evident at higher doses, at 50 µg/mL *C. microcoma* and *M. uncinata* reduced viability by 32 and 21%, and at 400 µg/mL proliferation was reduced by 71 and 82%, respectively.

**Table 3.** IC_50_ values of the plant extracts tested on the four cell lines. *P. angustifolium* stem demonstrates strong antiproliferative activity. N/A indicates that the extracts did not exhibit notable antiproliferative activity on the specific cell lines.

| Sample Name | Huh7 ($\mu\text{g/mL}$) | Hep3B ($\mu\text{g/mL}$) | Hep-55.1c ($\mu\text{g/mL}$) | A52 ($\mu\text{g/mL}$) |
| --- | --- | --- | --- | --- |
| *P. angustifolium* | 37.85 ± 1.2 | 79.18 ± 1.1 | 64.35 ± 3.1 | 80.21 ± 4.0 |
| *A. excelsa* | N/A | N/A | N/A | N/A |
| *C. microcoma* | 133.71 ± 6.3 | 94.29 ± 2.2 | 284.63 ± 14.2 | 182.27 ± 6.8 |
| *G. parviflora* | N/A | N/A | N/A | N/A |
| *M. uncinata* | 50.25 ± 3.1 | N/A | 134.19 ± 2.2 | 140.17 ± 2.9 |
| *G. australe* | N/A | N/A | N/A | N/A |
| *E. similis* | N/A | N/A | N/A | N/A |

**Figure 4.**
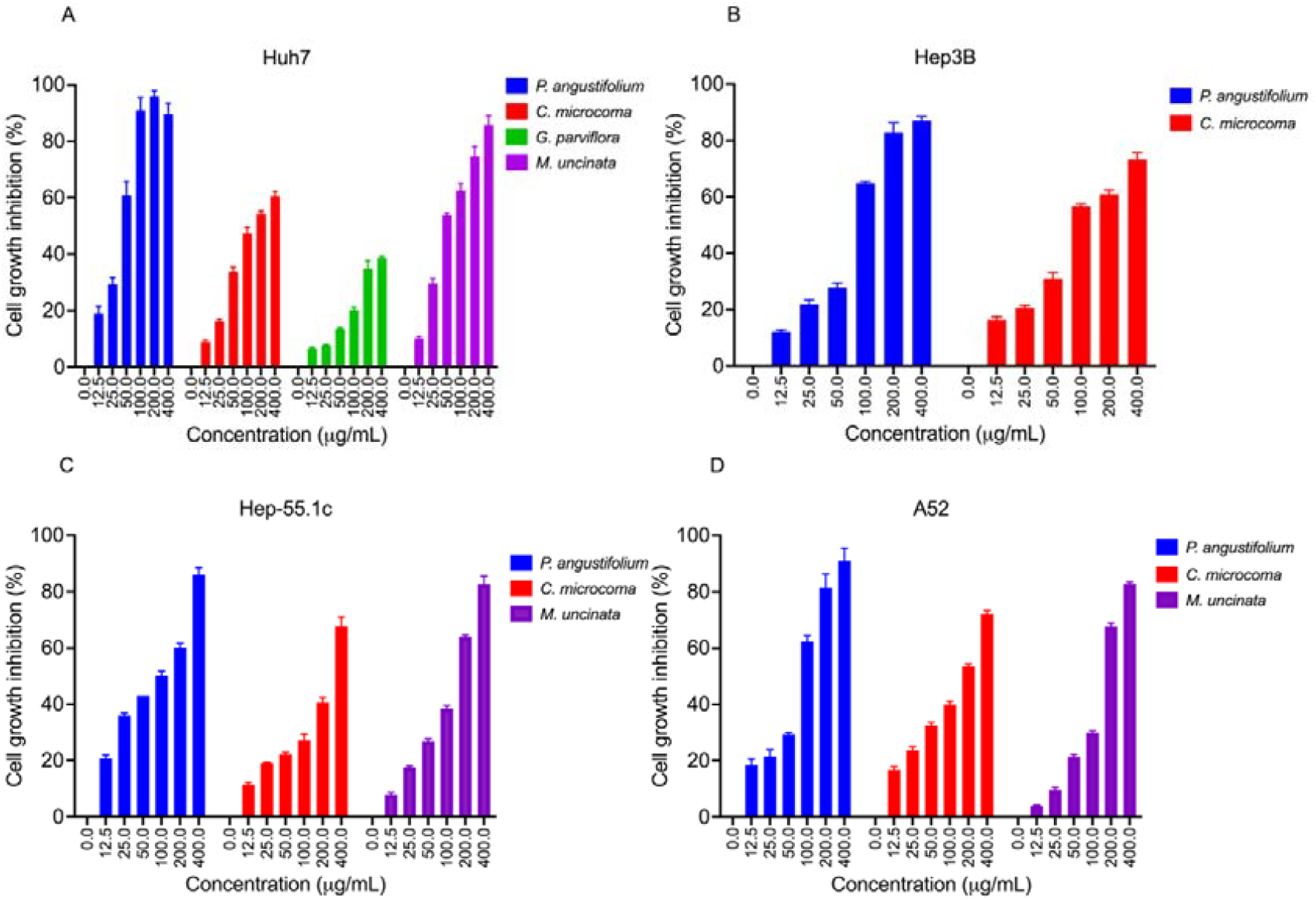
Concentration-dependent growth inhibition by Australian Indigenous medicinal plant extracts (12.5 to 400 μg/mL) on liver cancer cell lines Huh7, Hep3B, Hep-55.1c and A52. Cell viability was determined using the MTT assay, and growth inhibition (%) was calculated relative to untreated controls. Increased growth inhibition indicates stronger antiproliferative effects of the extracts. Data are expressed as mean ± SD (n = 3).

## 4.0 Discussion

The present study investigated Australian Indigenous medicinal plants customary used by the Iningai community for treating conditions associated with inflammation, pain, infection, and general illness, through integrated assessment of antioxidant, anti-inflammatory, and antiproliferative activities. The study targeted biological processes that are strongly linked to human hepatocellular carcinoma (HCC), namely oxidative stress, chronic inflammation, and uncontrolled cellular proliferation (Dharshini et al., 2023). Dysregulation of these processes contributes significantly to HCC progression through promotion of DNA damage, chronic inflammation, tumour survival, angiogenesis, and immune evasion. Plant extracts capable of modulating these pathways could lead to the development of novel HCC therapeutic agents. This study suggests that Australian Indigenous medicinal plants possess distinct biological activities, reinforcing the value of combining traditional knowledge with modern pharmacological evaluation. Importantly, the findings illustrate how Indigenous knowledge systems can guide rational prioritisation of medicinal species for biodiscovery, particularly where traditional applications align with experimentally observed biological activity.

Although Australian medicinal plants have increasingly attracted attention for their antioxidant and anti-inflammatory potential, relatively few studies have comprehensively integrated antioxidant, immunomodulatory, and anti-cancer screening. Previous work on Australian Indigenous medicinal plants has largely focused on antioxidant and antimicrobial activities, with limited investigation into their effects on inflammatory cytokines and liver cancer cell proliferation. A previous study revealed that eight Indigenous medicinal plants, scavenge free radicals by inhibiting DPPH with IC_50_ values ranging from 51.99 ± 1.17 to 2190.13 ± 2.16 μg/mL which were much lower compared to our extracts (8.87 ± 0.3 to 53.38 ± 3.7 μg/mL) (Akter et al., 2016). In another study, eight Iningai medicinal plants had antioxidant capacity with IC_50_ values ranging from 37.37 ± 1.01 to 206.50 ± 2.44 μg/mL and anti-inflammatory activity by modulating proinflammatory cytokines interferon-gamma (IFN-γ), interleukin 23 (IL-23), tumour necrosis factor (TNF), monocyte chemoattractant protein-1 (MCP-1) (Turpin et al., 2026). Therefore, our study represents one of the few integrated evaluations linking antioxidant capacity, inflammatory cytokine modulation, and antiproliferative activity in Australian Indigenous medicinal plants. Unlike previous studies focusing primarily on antioxidant or antimicrobial activity, the current work integrates tumour-relevant biological processes including oxidative stress, inflammation, and proliferation within a simple experimental framework.

The antioxidant assays demonstrated substantial variability amongst the extracts, with *C. microcoma* leaf and *G. parviflora* stem exhibiting the strongest DPPH radical scavenging and ferric-reducing activities, while *P. angustifolium* stem displayed comparatively weaker antioxidant capacity. These findings suggest the presence of redox-active phytochemicals. Previous phytochemical studies indicate that *G. parviflora* contains coumarins, alkaloids, flavonoids, and terpenoids (Dugan et al., 2024), which may contribute to the strong antioxidant activity observed here. Interestingly, several plants customarily used for treating pain, wounds, and inflammatory conditions showed strong antioxidant activity, partially supporting their ethnomedicinal use. However, a key finding of this study was that strong antioxidant activity did not reliably predict anti-inflammatory or anticancer efficacy. For example, *P. angustifolium* stem exhibited relatively weak antioxidant activity but demonstrated the strongest antiproliferative effects across all liver cancer cell lines tested. This highlights the limitation of relying only on chemical antioxidant assays when assessing therapeutic potential in complex biological systems.

The anti-inflammatory assay further demonstrated extract-specific immunomodulatory activity. *E. similis* leaf suppressed multiple pro-inflammatory cytokines, including IFN-α, TNF-α, IL-6, IL-12, IL-18, and IL-23, indicating strong anti-inflammatory potential. Interestingly, *E. similis* is traditionally used for fever reduction, pain relief, and respiratory conditions, which aligns with the observed cytokine suppression and provides pharmacological support for its customary application. In contrast, *G. parviflora*, traditionally used for pain and aches, demonstrated the strongest antioxidant activity, but only reduced IL-12. Whereas *P. angustifolium*, used for inflammatory and respiratory conditions, promoted TNF-α, IL-6, and inhibited IL-12, had the greatest antiproliferative effects despite moderate antioxidant activity. These observations suggest that ethnomedicinal applications have specific biological activities, although not uniformly across disease pathways. In contrast, *A. excelsa*, traditionally used as an antiseptic for wounds and skin infections, and *C. microcoma*, used for cuts, wounds and muscle pain, demonstrated selective cytokine modulation rather than broad suppression. *C. microcoma* leaf significantly reduced IL-12, IL-18, and IL-23, while *A. excelsa* reduced TNF-α and IL-18. Similarly, *M. uncinata*, traditionally used for respiratory and stomach ailments, preferentially suppressed IL-23, whereas *G. australe* stem reduced TNF-α but increased IL-18 levels. These findings suggest that individual extracts may influence distinct inflammatory pathways, reflecting both their phytochemical complexity and their diverse traditional medicinal applications. Importantly, *E. similis* leaf did not demonstrate antiproliferative activity in liver cancer cells, indicating targeted immunomodulation rather than induction of cellular stress or cytotoxicity.

Our study also showed limited concordance between anti-inflammatory activity and antiproliferative effects. Extracts such as *E. similis* which demonstrated strong cytokine suppression were not cytotoxic against liver cancer cells. Conversely, *P. angustifolium* stem demonstrated strong antiproliferative activity despite exhibiting weak antioxidant and antiinflammatory effects. This could be attributed to the biological differences between immune cells and cancer cells. Macrophages adapt to tolerate inflammatory and oxidative stress during immune responses, whereas HCC cells already exist under elevated oxidative and metabolic stress conditions. Consequently, compounds capable of inducing additional mitochondrial or oxidative stress may selectively impair cancer cell survival and exert limited anti-inflammatory activity. This suggests that anticancer efficacy may depend more on the ability of extracts to disrupt tumour cell stress-response pathways than their antioxidant capacity.

Our findings reinforce the importance of context-dependent evaluation of natural products. Plant extracts are inherently complex, containing multiple bioactive constituents that engage distinct molecular targets across different cell types (Dar et al., 2023). By integrating antioxidant assays, cytokine profiling, and cancer cell viability analyses, we demonstrate that bioactivity emerges from the interaction between extract phytochemical composition and cellular context, rather than from isolated molecular properties. Our findings help scientifically confirm the efficacy of Indigenous medicinal knowledge and highlight Australian native flora as an underexplored source of selective therapeutic leads.

Despite these promising findings, several limitations should be acknowledged. The present study utilised crude plant extracts. Therefore, the specific compounds responsible for the observed activities remain unidentified. Moreover, while antioxidant, cytokine, and proliferation assays provide valuable insights, they do not fully replicate tumour microenvironments. Future work involving phytochemical isolation, apoptosis analysis, ROS modulation assays, and *in vivo* validation will be essential to confirm therapeutic relevance and elucidate mechanisms of action. Such studies may contribute to the development of selective anticancer agents derived from Australian Indigenous medicinal plants.

## 5.0 Conclusion and future directions

In summary, this study reveals that Australian Indigenous medicinal plants exhibit distinct antioxidant, immunomodulatory, and anticancer profiles, rather than uniform bioactivity across systems. These findings position Australian Indigenous medicinal plants as promising yet underexplored sources of selective bioactive compounds and reinforce the value of Indigenous knowledge-guided biodiscovery approaches for anticancer drug discovery.

While the present study provides strong functional insights, further mechanistic investigations are warranted to delineate the molecular pathways underpinning the observed effects. Future studies incorporating apoptosis markers, ROS modulation assays, cell-cycle analysis, and pathway-specific reporter systems would strengthen the study’s interpretation. Additionally, phytochemical profiling and bioassay-guided isolation will be essential to identify the constituents responsible for the divergent bioactivities observed. In summary, these future studies may contribute to the development of novel, selective anticancer agents derived from Australian Indigenous medicinal plants.

## Author contributions CRediT

**Felix Wambua Muema:** Conceptualization, Data curation, Methodology, Formal analysis, Writing – original draft, Writing – review & editing. **Suzanne Thompson:** Conceptualization, Data curation, Writing – review & editing. **Gerry Turpin:** Conceptualization, Data curation, Writing – review & editing. **Graham Ambridge:** Conceptualization, Data curation, Writing – review & editing. **Joanne Jamie:** Conceptualization, Data curation, Writing – review & editing. **Darren Crayn:** Conceptualization, Data curation, Writing – review & editing. **Catherine M. Miller:** Conceptualization, Data curation, Formal analysis, Methodology, Project administration, Resources. **Lionel Hebbard:** Conceptualization, Data curation, Formal analysis, Methodology, Project administration, Funding acquisition. **Phurpa Wangchuk:** Conceptualization, Data curation, Formal analysis, Methodology, Project administration, Funding acquisition.

## Acknowledgement

The authors gratefully acknowledge the Yambangku Aboriginal Cultural Heritage & Tourism Development Aboriginal Corporation (YACHATDAC) for generously sharing their medicinal plant knowledge and for their contribution to the collection and provision of plant materials from Turrabura Station on Iningai Country. We recognise and deeply value the Indigenous knowledge that informed this research and acknowledge the continuing connection of the Iningai people to their Country, culture, and traditional knowledge.

This research was conducted at the Cairns (Nguma-bada) Campus of James Cook University. We acknowledge the Traditional Custodians of the lands on which the plant materials were collected, and the research was undertaken, and pay our respects to their Elders past and present. We acknowledge Australian Institute of Tropical Health, and Medicine (AITHM), Cairns Campus, James Cook University for access to its instrumentation and Tenzin Jamtsho (JCU) for his help in BD LSRFortessa analysis.

